# VIGAT: VLM-guided gene activity quantification from single-cell ATAC-seq

**DOI:** 10.64898/2026.10.16.759698

**Authors:** Beijie Ji, Zhicheng Ji

**Affiliations:** Department of Biostatistics and Bioinformatics, Duke University School of Medicine, Durham, NC, USA; Pierre R. Lamond Department of Electrical and Computer Engineering, Duke University Pratt School of Engineering, Durham, NC, USA

## Abstract

**Summary:** Gene activity quantification from single-cell ATAC-seq is challenging because genomic regions informative of cell identity can vary across genes and tissues, while broadly accessible regions may provide limited cell-type-discriminative information. We developed VIGAT, a vision-language model (VLM)-guided tool that automates visual interpretation of ATAC coverage plots for adaptive gene activity quantification. VIGAT jointly examines accessibility patterns across multiple marker genes, integrating genomic annotations and biological context to select gene-specific regions enriched for cluster-discriminative signals. Selected regions are used to generate a gene-by-cell activity matrix. Across 196 paired single-cell RNA-seq and ATAC-seq datasets, VIGAT outperformed five existing methods in recovering marker-associated accessibility enriched in the corresponding cell populations, demonstrating the utility of VLM-guided visual reasoning for scalable gene activity quantification.

**Availability and implementation:** VIGAT is implemented as an open-source R package and is freely available at https://github.com/beijie02/VIGAT.

## Introduction

Single-cell ATAC-seq profiles chromatin accessibility at single-cell resolution^1^. Gene activity quantification aggregates accessibility signals from genomic regions near or within genes, such as promoters and gene bodies, to estimate gene-level activity for downstream cell type characterization and biological interpretation. However, within these genomic regions, some segments exhibit cell-type-specific accessibility, whereas others remain broadly accessible across cell populations and provide limited information for distinguishing cell types^2–4^. Furthermore, the locations of cell-type-discriminative accessibility signals can vary across genes and tissue types. These characteristics highlight the importance of selecting informative genomic regions adaptively, rather than applying the same predefined rules across genes and datasets.

Existing gene activity quantification methods generally rely on predefined genomic regions, distance-based weighting schemes, or inferred regulatory connections. For example, Signac^5^ and SnapATAC2^6^ quantify accessibility within gene bodies and fixed upstream extensions, while ArchR^7^ and MAESTRO^8^ incorporate distance-based weighting and genomic annotations. Cicero^9^ infers regulatory connections through chromatin coaccessibility. However, these methods do not explicitly adapt genomic region selection to prioritize cell-type-discriminative accessibility signals. Such informative regions can often be identified by visually inspecting ATAC coverage plots, which reveal accessibility differences across cell clusters alongside gene annotations and called peaks. However, manual inspection and selection of genomic regions across large numbers of genes and datasets is labor-intensive and difficult to scale.

Here, we present VIGAT, a vision-language model (VLM)-guided tool that automates visual inspection of ATAC coverage plots to adaptively identify informative genomic regions for gene activity quantification. VIGAT jointly presents coverage plots for multiple marker genes to a VLM, enabling it to interpret accessibility differences across cell clusters in the context of gene identities, genomic annotations, and biological knowledge to identify regions enriched for cell-type-discriminative accessibility while avoiding broadly shared signals. The selected regions are then used to quantify ATAC fragments and construct a gene-by-cell activity matrix, without requiring explicit cell type annotations. Across 196 paired single-cell RNA-seq and ATAC-seq datasets, VIGAT outperformed five existing methods in capturing marker-associated accessibility enriched in the corresponding cell populations.

## Materials and methods

### VIGAT software implementation

VIGAT takes as input a processed Seurat object and a user-provided list of marker genes. The Seurat object can be generated using the standard Signac workflow and should contain a `ChromatinAssay` with ATAC fragments, called peaks, and gene annotations, together with cell cluster assignments. No matched RNA-seq data or cell type annotations are required.

For each marker gene, VIGAT generates a coverage plot spanning 5 kb upstream and downstream of the transcription start site (TSS) using Signac `CoveragePlot()`, displaying ATAC accessibility across clusters alongside gene annotations and called peaks. By default, clusters containing fewer than 50 cells are excluded, and at least two clusters are required. The coverage plots are jointly submitted to the OpenAI Responses API together with gene names, strand orientations, tissue and species information, and called-peak coordinates. Guided by a standardized prompt (Supplementary Information), the VLM selects one compact genomic range per gene, generally at least 1 kb long, prioritizing promoter-proximal, cluster-discriminative accessibility signals while avoiding broadly shared signals and regions associated with neighboring genes. We used GPT-5.5 in this study, although other models can be specified through the `openai_model` argument. The selected coordinates are converted into `GRanges` objects, and overlapping ATAC fragments are quantified for each cell using Signac `FeatureMatrix()`. VIGAT outputs the selected genomic ranges, their coordinates in tabular format, and a sparse gene-by-cell raw count matrix, with optional coverage plots highlighting the selected regions. VIGAT was implemented in R 4.4.3 using Seurat 5.5.1, Signac 1.17.1, and GenomicRanges 1.58.0.

### Benchmark and evaluation

We evaluated VIGAT using 196 paired single-cell RNA-seq and ATAC-seq datasets from CompassDB^10^, in which both modalities were measured in the same cells and cell type annotations were derived exclusively from the RNA-seq data. For each dataset, ATAC cells were clustered using the standard Signac workflow^5^, including TF-IDF normalization, latent semantic indexing (LSI), and graph-based clustering with the smart local moving algorithm. Five clustering resolutions (0.1–0.5) were evaluated, and the resolution with the lowest mean mismatch rate between ATAC clusters and RNA-derived cell type annotations was selected. Specifically, each ATAC cluster was assigned the majority RNA cell type of its constituent cells, and the mismatch rate was calculated as the proportion of cells whose RNA annotations differed from the assigned cluster label, averaged equally across clusters. At the selected resolution, these majority-vote labels served as reference cell types for evaluation.

Datasets were filtered to exclude ATAC clusters with fewer than 50 cells, require at least two distinct reference cell types, and ensure that the second most abundant RNA cell type contained more than 50 cells. Marker genes were obtained from DISCO^11^, and clusters without valid marker gene mappings were excluded, with at least two reference cell types required to remain. These criteria yielded 196 datasets. RNA-derived cell type annotations were used for benchmark preparation and evaluation but were not provided to VIGAT during genomic range selection, for which ATAC clusters were identified only by number. VIGAT was compared with Signac, SnapATAC2, ArchR, Cicero, and MAESTRO using their documented workflows, with all methods evaluated on common cells and marker genes.

For each marker gene *g* associated with reference cell type *c*, we calculated the proportion of cells with nonzero gene accessibility in the target cell type, *p*_*g,c*_, and in pooled non-target cell types, *p*_*g,−c*_. The relative marker ATAC detection score was defined as *R*_*g,c*_ = *p*_*g,c*_*/p*_*g,−c*_, with zero denominators replaced by the smallest positive pooled non-target detection proportion for that method across the benchmark. Scores were averaged across marker genes within each cell type, then equally across reference cell types within each dataset, and finally across datasets. Higher scores indicate stronger enrichment of marker-associated accessibility in the corresponding cell populations.

## Results

We first illustrated VIGAT’s genomic range selection using a representative human islet dataset containing seven reference cell types and 15 marker genes (Figure 1a–c). All 15 marker genes and their corresponding ATAC coverage plots were jointly provided to GPT-5.5 for genomic range selection. For visualization, Figure 1a–c presents two representative marker genes, *TTR* and *SPP1*, associated with alpha cells and TUBA1A ductal cells, respectively^11^. The VLM interpreted accessibility patterns across ATAC clusters together with gene identities, genomic annotations, and called peaks to select one informative genomic range per gene (Figure 1a). As illustrated in Figure 1b,c, the selected ranges captured localized accessibility signals enriched in the corresponding cell populations while excluding nearby regions with less discriminative accessibility patterns. Cell type identities were not provided to the VLM and were added to the plots only for visualization.

**Figure 1.**
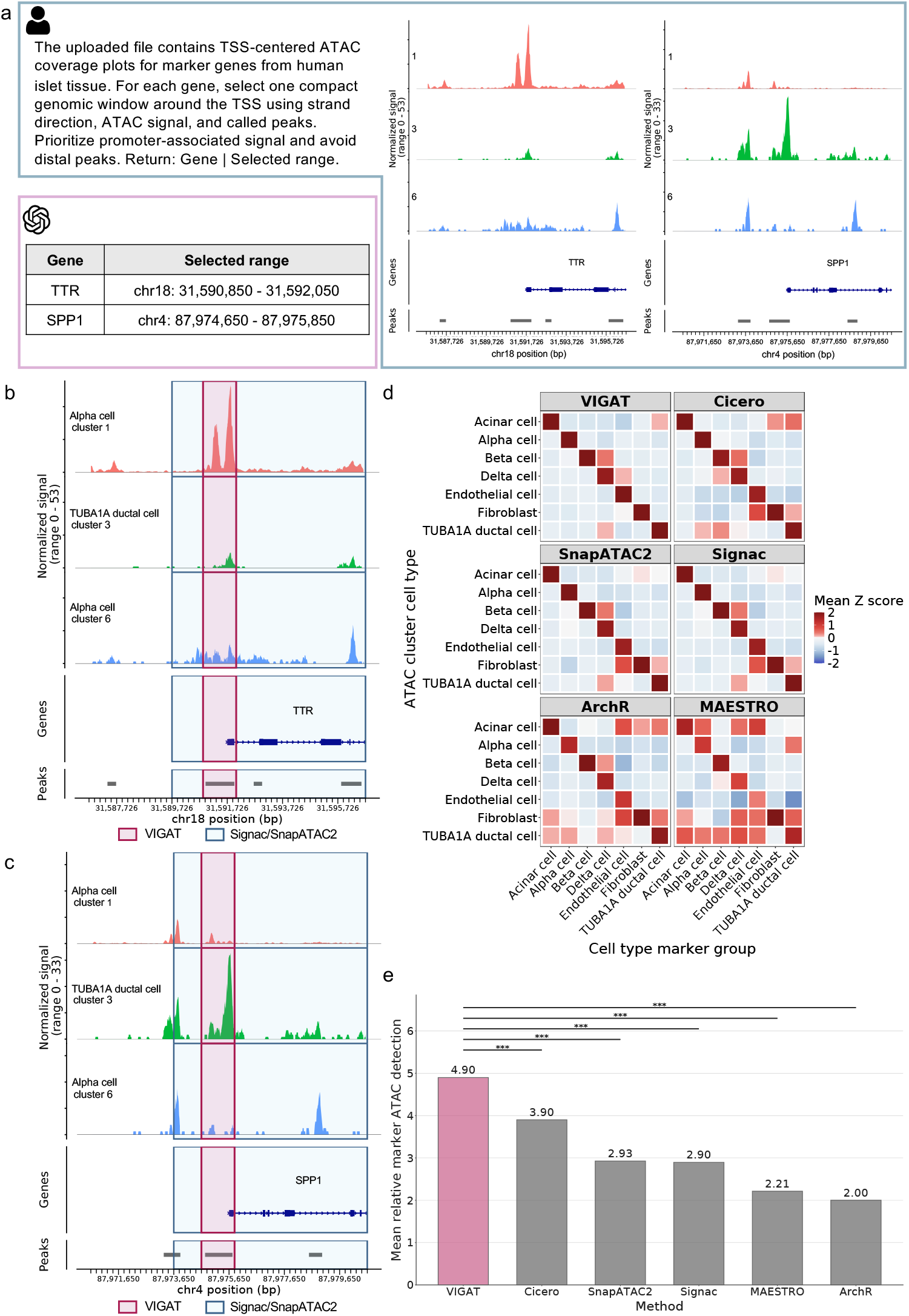
VIGAT workflow and benchmark performance. **a**, Illustration of VLM-guided range selection using two representative marker genes, *TTR* and *SPP1*, from 15 marker genes jointly analyzed in a human islet dataset. The VLM integrates ATAC coverage patterns, gene identities, annotations, and called peaks to select informative genomic ranges. **b-c**, ATAC coverage profiles for *TTR* (**b**) and *SPP1* (**c**) across three representative ATAC clusters, highlighting VIGAT-selected regions and the corresponding regions used by Signac and SnapATAC2 within the displayed windows. RNA-derived reference cell type identities are shown for interpretation but were withheld from the VLM. **d**, Comparison of cell-type-specific marker accessibility across six methods using all 15 marker genes. Detection proportions were standardized as Z-scores across cell types and averaged within marker groups to generate heatmaps. Higher diagonal values indicate stronger enrichment in the corresponding cell types. **e**, Mean relative marker ATAC detection scores across 196 paired single-cell RNA-seq and ATAC-seq datasets. Higher scores indicate greater marker accessibility enrichment in the corresponding cell types. Statistical significance was assessed using paired two-sided Wilcoxon signed-rank tests with Benjamini–Hochberg correction. *** indicates adjusted *P <* 0.001.

We next evaluated cell-type-specific marker accessibility across the six methods using all 15 marker genes in the same islet dataset (Figure 1d). Using the same marker ATAC detection proportions as in the overall benchmark, we visualized cell-type-specific accessibility patterns in a cell type-by-marker group heatmap. VIGAT exhibited the strongest overall diagonal enrichment pattern, indicating that its selected genomic ranges more effectively captured marker-associated accessibility in the corresponding cell populations. In comparison, competing methods showed weaker enrichment in the expected cell types or increased accessibility in non-target populations for several marker groups.

Finally, we benchmarked VIGAT against five existing gene activity quantification methods across 196 paired single-cell RNA-seq and ATAC-seq datasets from CompassDB (Figure 1e). Performance was evaluated using the relative marker ATAC detection score described in Methods, which measures the enrichment of marker-associated accessibility in the corresponding reference cell type relative to other cell types. VIGAT achieved the highest mean score of 4.90, outperforming Cicero (3.90), SnapATAC2 (2.93), Signac (2.90), MAESTRO (2.21), and ArchR (2.00), representing a 25.6% improvement over the second-best method, Cicero. These results demonstrate that VLM-guided selection of informative genomic regions more effectively captures cell-type-enriched accessibility signals across diverse datasets.

## Conclusion

VIGAT enables scalable, adaptive gene activity quantification by leveraging VLMs to identify informative genomic regions from ATAC coverage plots, without requiring cell type annotations. Its superior performance across diverse datasets demonstrates the potential of VLM-guided visual reasoning for improving gene activity quantification in single-cell ATAC-seq analysis.

## Acknowledgments

This work was supported by the National Institutes of Health under award numbers U54AG075936, R35GM154865, and U54AI191253.

## Author contributions

Z.J. conceived the study. B.J. developed the method and software and performed the analyses. B.J. and Z.J. wrote the manuscript.

## Data availability

The benchmark datasets used in this study are publicly available from CompassDB^10^.

