## Supplementary Information for "VIGAT: VLM-guided gene activity quantification from single-cell ATAC-seq"

### Prompt used by VIGAT

The following prompt was used by VIGAT to select gene-specific genomic windows from TSS-centered ATAC coverage plots. Text enclosed in angle brackets indicates information that was automatically populated by the software for each analysis.

```
The uploaded PDF file(s) contain TSS-centered ATAC CoveragePlots for marker genes. For each
marker gene plot, select one compact absolute genomic window. The selected window should be
suitable for calculating the ATAC signal of that marker gene later.

Context: ATAC data from <SPECIES> <TISSUE>. Cluster identities are unknown. Use the ATAC
coverage signal, gene model, genomic coordinates, marker gene name, TSS direction provided
after each gene name, and called Peaks if available.

Markers:
<MARKERS>

Called Peaks:
<CALLED_PEAKS>

Instructions:

1. First, identify the target marker gene for each coverage plot.

2. Locate the marker gene's TSS/promoter region based on its strand orientation. For a "+"
strand gene, the promoter region lies toward decreasing genomic coordinates. For a "-"
strand gene, the promoter region lies toward increasing genomic coordinates.

3. Choose one compact window near the marker gene's TSS/promoter region.
    - If a called Peak overlaps with the promoter signal, prioritize a window that overlaps
    this Peak and captures the clearest cluster-specific ATAC signal. The window does not
    need to span the entire called Peak if only a subset of the region is informative. You
    may include a small, contiguous neighboring region if it helps resolve cluster
    differences.
    - If no called Peak overlaps the promoter signal but a clear, cluster-specific coverage
    signal is visible near the TSS, define a compact window based on the coverage profile
    alone.

4. Do not select a window solely based on the highest signal intensity. Avoid distal peaks
far from the promoter, broad non-specific signals shared across many clusters, and signals
originating from adjacent genes.

5. Use a minimum window size of 1 kb by default. A window smaller than 1 kb should only be
used if the promoter-side Peak or coverage signal is exceptionally sharp and well-isolated.
Otherwise, expand the window modestly around the informative signal while avoiding unrelated
neighboring noise.

6. Before final output, verify that each selected start and end coordinate is within the
displayed x-axis range for that gene.

Output Format:

First, provide one row per gene:
Gene; TSS direction; chr:start-end; Size; Anchor evidence (called Peak(s) or coverage-only);
Strongest useful clusters; Brief rationale.

Then, provide the selected windows as R code for only the genes with a valid selected range.
Keep the same gene order as in the row-format output, and omit genes whose range is NA or
cannot be determined.

vigat_windows <- GRanges(
  seqnames = c(
```

```
    "chr1", "chr2"
  ),
  ranges = IRanges(
    start = c(
      100000, 200000
    ),
    end = c(
      101000, 201000
    )
  ),
  gene = c(
    "GENE1", "GENE2"
  )
)

============================================================
```

Here, <SPECIES> and <TISSUE> were replaced by the species and tissue supplied by the user. <MARKERS> was automatically populated with the marker genes and their TSS directions, and <CALLED_PEAKS> was populated with the called ATAC peaks associated with the input data when available.
